# Automated extraction of primary cilia-based biomarkers reveals ageing of cells

**DOI:** 10.64898/2026.04.14.718373

**Authors:** Juan Esteban Montes-Montoya, Zoe Tryfonos, Ji Eun Lee, Hyuk Wan Ko, Soo-Hyun Kim, Constantino Carlos Reyes-Aldasoro

## Abstract

We present an automated image-analysis methodology for quantitative assessment of primary cilia in cultured human primary fibroblasts. The proposed approach implements a modular processing pipeline combining deep-learning based nuclear segmentation with intensity-driven segmentation of the ciliary axoneme and basal body. These partial segmentations are integrated using a distance-based association strategy, enabling automated reconstruction of individual cilia and subsequent extraction of geometrical features. Performance was evaluated against manually segmented ground truth. While manual annotations showed a systematic underestimation of cilia length, both automated and manual analyses produced consistent population-level trends and identical statistical discrimination across sample groups. Application of the pipeline revealed a progressive reduction in cilia length and ciliation frequency with increasing passage number, demonstrating the sensitivity of the method to subtle morphological changes. The proposed framework enables scalable, reproducible, and objective quantification of primary cilia morphology and provides a computational tool applicable to high-throughput studies of cellular ageing and related phenotypes.

All the code related to this work is available through GitHub: https://github.com/reyesaldasoro/Cilia.

## 1 Introduction

The primary cilium is a microtubule-based organelle, which serves as a critical signalling hub that governs key cellular functions, including intercellular communication, protein trafficking, secretion, recycling, and genome stability [10].

Notably, ciliary alterations are closely linked to ageing. For instance, cells from patients with Progeria (a childhood-onset disorder characterised by accelerated ageing) exhibit significantly shortened primary cilia when compared to healthy controls [5]. Alternations of primary cilia have also been observed in human cells undergoing senescence when cells have stopped dividing in tissue culture [1, 4]. Significant evidence supports a critical role of primary cilia in the pathogenesis of a wide range of common ageing-associated diseases [10, 12, 7, 3]. These data collectively suggest that ageing associates with altered function or structure of primary cilia and healthy primary cilia are essential for longevity and cellular resilience. However, it remains unclear whether ciliary morphology changes can serve as robust and quantifiable biomarkers of healthy ageing.

Microscopic imaging after staining with cilia-specific fluorescent markers, measuring cilia length, number, and morphology, is the gold standard for assessing cilia status [6]. Current approaches are limited by low-throughput imaging workflows, reliance on manual or semi-manual annotation, and analysis focused on a small number of features. These limitations reduce reproducibility and make large-scale analysis of cells and cilia difficult. Approaches based on computer vision and machine-learning can offer a route to standardised cilia analysis by reducing human bias and enabling accurate, reproducible, and scalable quantification of ciliary features from florescence microscopy images.

In this study, we introduce a fully automated computational framework for the quantitative analysis of primary cilia from fluorescence confocal microscopy images. The proposed method performs end-to-end segmentation of individual cells, cilia axonemes and basal bodies without user intervention, enabling objective and reproducible feature extraction. The pipeline was validated against manual measurements, demonstrating high accuracy and robustness in estimating ciliary geometry. We then applied the methodology to immunofluorescence images of human primary fibroblasts across replicative ageing to quantify age-dependent morphological changes using cilia-derived features. By improving the scalability, consistency, and throughput of cilia segmentation and measurement, and assessing the feasibility of intensity-based segmentation for cilia image analysis, this framework provides an image-analysis tool suitable for large-scale studies and supports the systematic evaluation of ciliary morphology as a biomarker of cellular ageing and age-associated disease.

## 2 Materials and Methods

### 2.1 Materials

#### Cell Culture

Primary neonatal foreskin fibroblasts were kindly provided by Dorothy Bennett [13]. Cells were recovered at passage 13 (P13) and subsequently expanded to passages P16, P22 and P28 to model replicative ageing. Dulbecco’s Modified Eagle Medium (DMEM, Sigma-Aldrich) supplemented with 10 % fetal bovine serum (FBS, Sigma-Aldrich), 1 % L-glutamine (Sigma-Aldrich), and 1 % penicillin-streptomycin (Sigma-Aldrich) was used, with regular media changes every 3-4 days. Cultures were maintained at 37 °C in a humidified atmosphere containing 5% CO_2_.

#### Pre-staining Culture Conditions

Fibroblasts were seeded onto 13 mm round glass coverslips coated with 0.01 % Poly-D-Lysine (Gibco) at a density of 3×10^4^ cells per well in serum-containing medium. To induce ciliogenesis, cells were serum-starved for 48 hours prior to fixing.

#### Antibodies

Primary antibodies used for immunofluorescence staining were anti-ARL13B (Proteintech, cat. no. [17711-1-AP], diluted at 1:4000), and mouse monoclonal anti-*γ*-tubulin (Sigma-Aldrich, cat. no. [T6557], diluted at 1:600). Secondary antibodies were Alexa Fluor 555 cross-absorbed goat anti-mouse (Invitrogen, cat. no. [A-21422], diluted at 1:5500) and Alexa Fluor 488 cross-absorbed goat anti-rabbit (Invitrogen, cat. no. [A-11008], diluted at 1:5500).

#### Immunofluorescence

Coverslips containing adherent cells were washed once in Phosphate Buffered Saline (PBS, Dulbecco) and fixed in ice-cold 4 % formaldehyde (ThermoFisher) for 10 minutes. Following fixing, samples were washed three times with PBS and permeabilised in PBS containing 0.2% Triton X-100 (Sigma-Aldrich) for 10 minutes. Cells were subsequently washed three times with PBS and blocked for 1 hour at room temperature in 10 % heat-inactivated goat serum prepared in PBS containing 0.2% Triton X-100 (the blocking buffer). This was followed by overnight incubation with primary antibodies at 4°C. After primary antibody incubation, samples were washed three times with PBS and incubated with secondary antibodies for 1 hour at room temperature in the dark. Following three final washes with PBS, coverslips were mounted using Vectashiled antifade medium containing DAPI (Vector Labs).

#### Confocal Image Acquisition

Images were acquired using a Nikon A1plus confocal microscope mounted on a Ti2 platform, equipped with a 60X Apo oil-immersion objective (numerical aperture 1.4; refractive index 1.515). The image resolution was set to 1024 × 1024 pixels, corresponding to a spatial calibration of 0.21 μm /pixel. Three fluorescence channels were collected using excitation wavelengths of 405 nm, 488 nm, and 561 nm, with corresponding emission wavelengths of 450 nm, 525 nm, and 595 nm. Z-stacks consisting of 10 optical planes were acquired using identical setting across all samples.

### 2.2 Methods

#### Segmentation Pipeline

The fluorescent channels of the images were analysed separately. The blue channel contained the DAPI-stained nuclei (Fig. 2a). These were segmented with the generalist segmentation algorithm *cellpose* [11]. Cellpose is a deep-learning based algorithm, which has been pre-trained for a variety of image types with over 70,000 segmented objects and has been validated for single-cell segmentation [9]. Nuclei segmentation with cellpose was satisfactory without need of training or fine-tuning with images related to the cilia dataset (Fig. 2b). A few parameters were easily determined: cell diameter, flow error threshold (acceptable error between the flow fields output by the network and flow fields reconstructed from the predicted label), cell probability threshold (balance between recall of cells against the more accurate boundaries of the detected cells). Watersheds were calculated from Euclidean distance maps (Fig. 2c) as a tool to isolate individual cells and to allocate cilia axoneme and basal bodies to each individual cell (Fig. 2d).

**Fig. 1.**
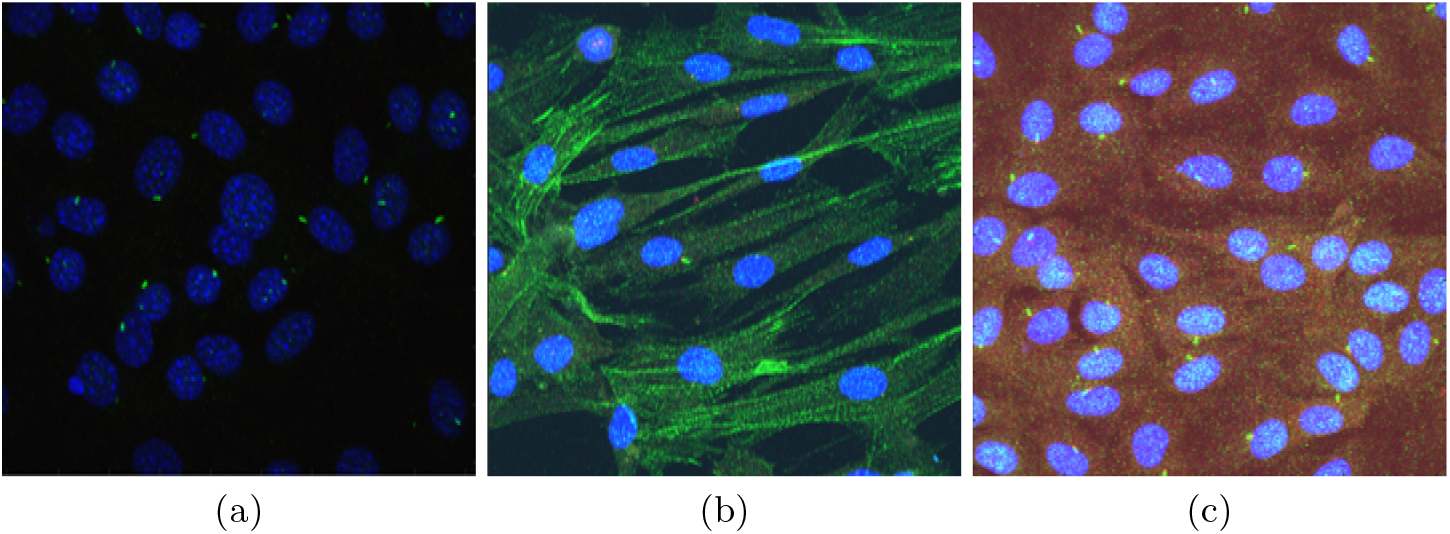
Representative examples of the cilia datasets. Human primary fibroblasts were serum-starved for 24-48 hrs. Nuclei were counterstained with DAPI (blue), the ciliary axoneme was labelled with anti-ARL13B (green), and the basal body was labelled with anti-*γ*-tubulin (red). The images illustrate variability in cell and structure intensities and background signal across samples: (a) predominantly ciliary signal observed in the green channel, (b) over-exposure in green channel showing signals from cilia and fibre-like structures, and (c) elevated background signal in the green and red channel.

**Fig. 2.**
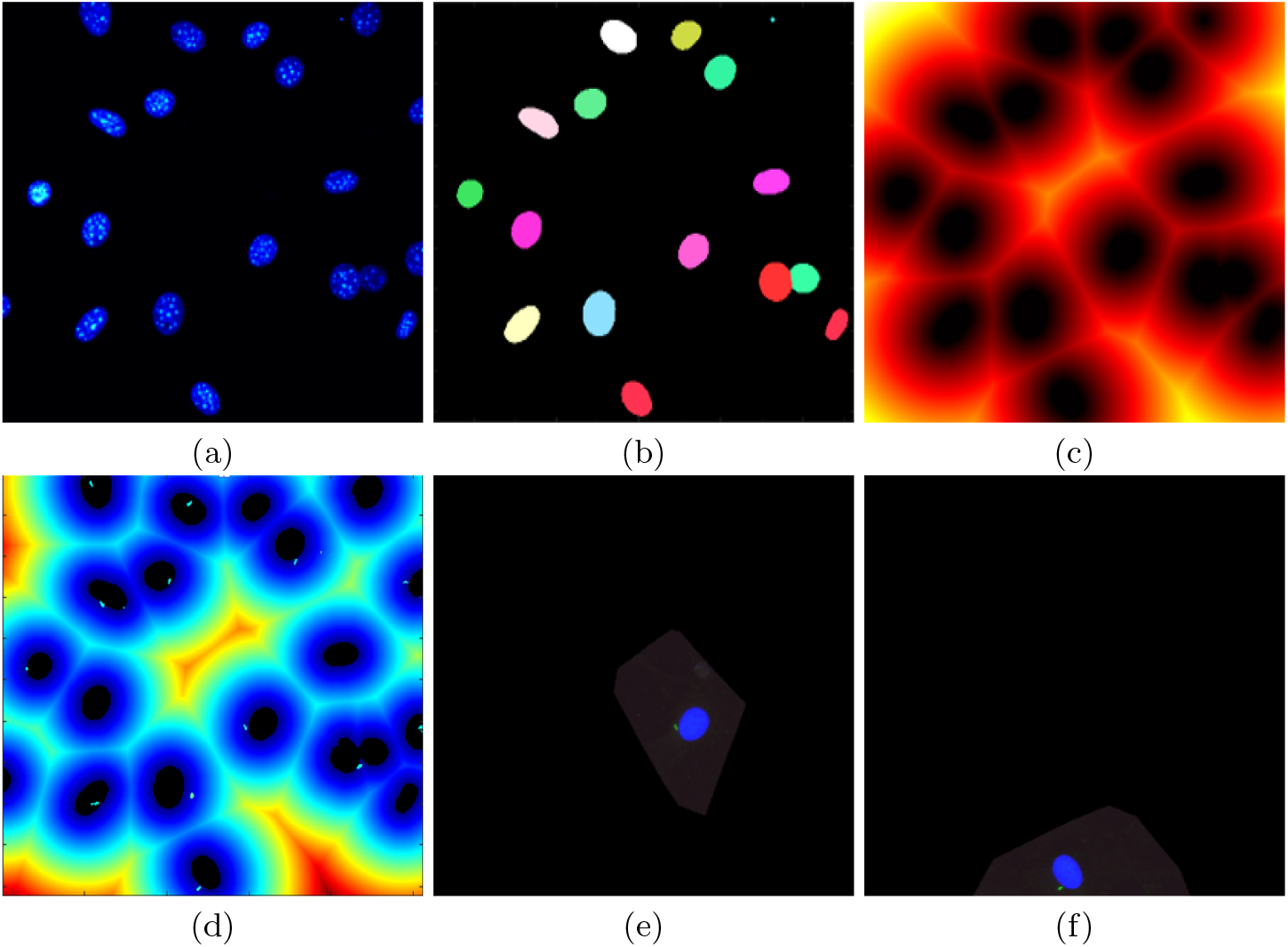
Illustration of the nuclei segmentation. (a) Blue channel containing nuclei. (b) Instance segmentation obtained with cellpose; it can be noticed that two overlapping nuclei were correctly segmented (bottom right). (c) Distance map based the nuclei segmentation, this map is used to isolate regions corresponding to individual cells. (d) Segmented cilia overlaid in distance map; this illustrates how cilia can be allocated to the corresponding nuclei and artifacts can be detected. (e,f) Two examples of individual cells and their cilia.

The green channel contained the intensity of the cilia axoneme and the red channel contained the information related to the basal bodies. These two were analysed for fluorescence intensity peaks in a strategy similar to that proposed in [8]. Each image was analysed in two separate directions (rows/columns) and for each, intensity peaks were detected (Fig. 3a,b). These peaks were considered to be a detections when intensity peaks co-located in both directions. This strategy, combined with a pre-determined peak prominence and distance between peaks removed peaks that could be caused by noise or very low intensities. Then, the peaks of both channels (Fig. 3c) were combined and cilia detections were only valid when the peaks in the green channel (cilia axoneme), co-located with peaks in the red channel (basal body) (Fig. 3d).

**Fig. 3.**
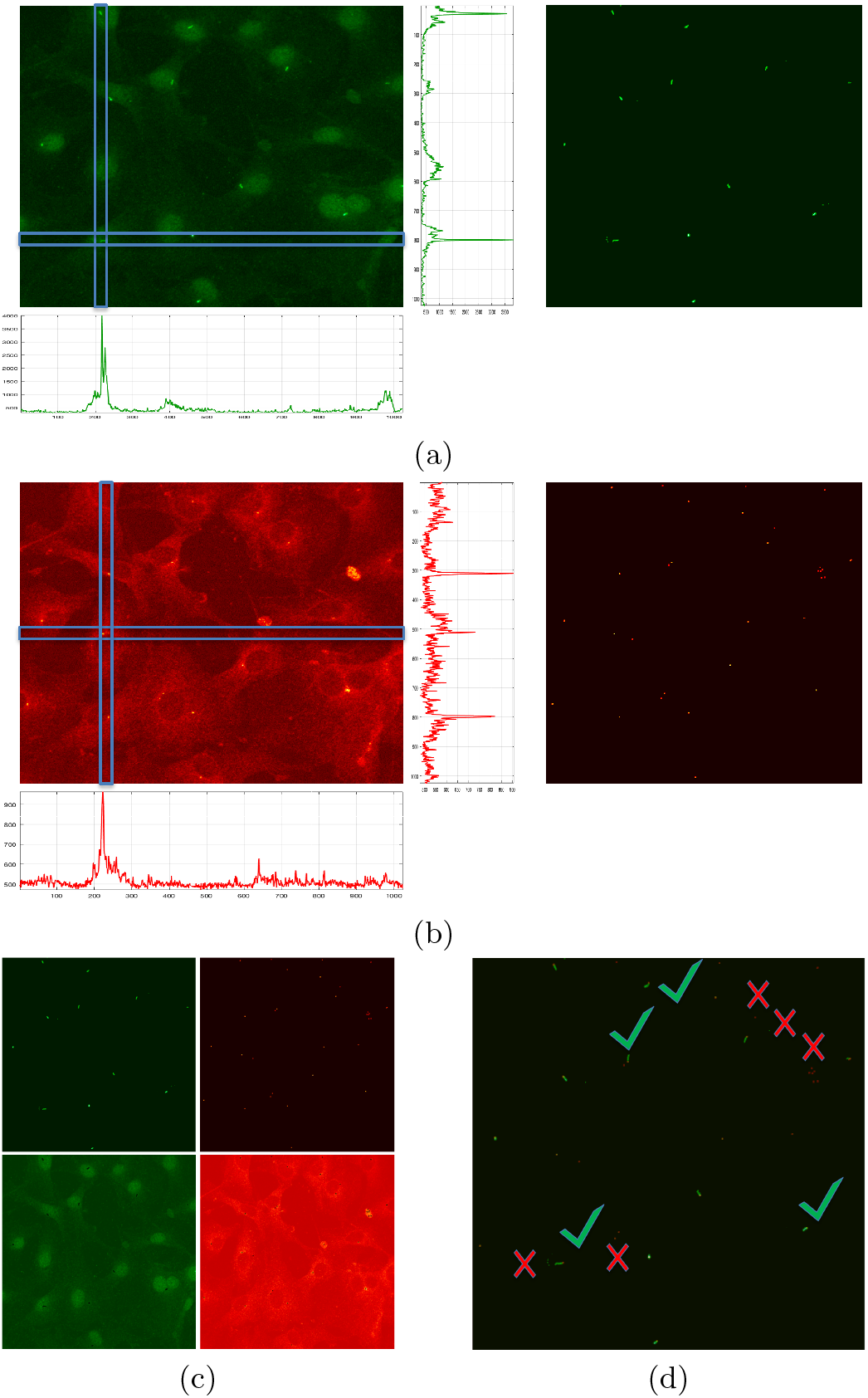
Intensity-based segmentation of (a) cilia in the green channel, and (b) basal body in red channel. In both cases, individual columns and rows of the images were scanned to detect peaks of intensities. These peaks were considered to be a detections when intensity peaks co-located in both directions. Then, the peaks of both channels (c) were combined, and detections in the green channel that co-located with detections in the red channel were considered as cilia (green ticks in (d)), or discarded otherwise (red crosses in (d)). To aid visualisation, green and red channels with the peaks removed are included in (c).

The output of the processing of the individual channels was combined to provide a final per-cell segmentation (Fig. 4). The pipeline was tested with a variety of cells with very different characteristics, and satisfactory results were obtained for all cases (Fig. 5).

**Fig. 4.**
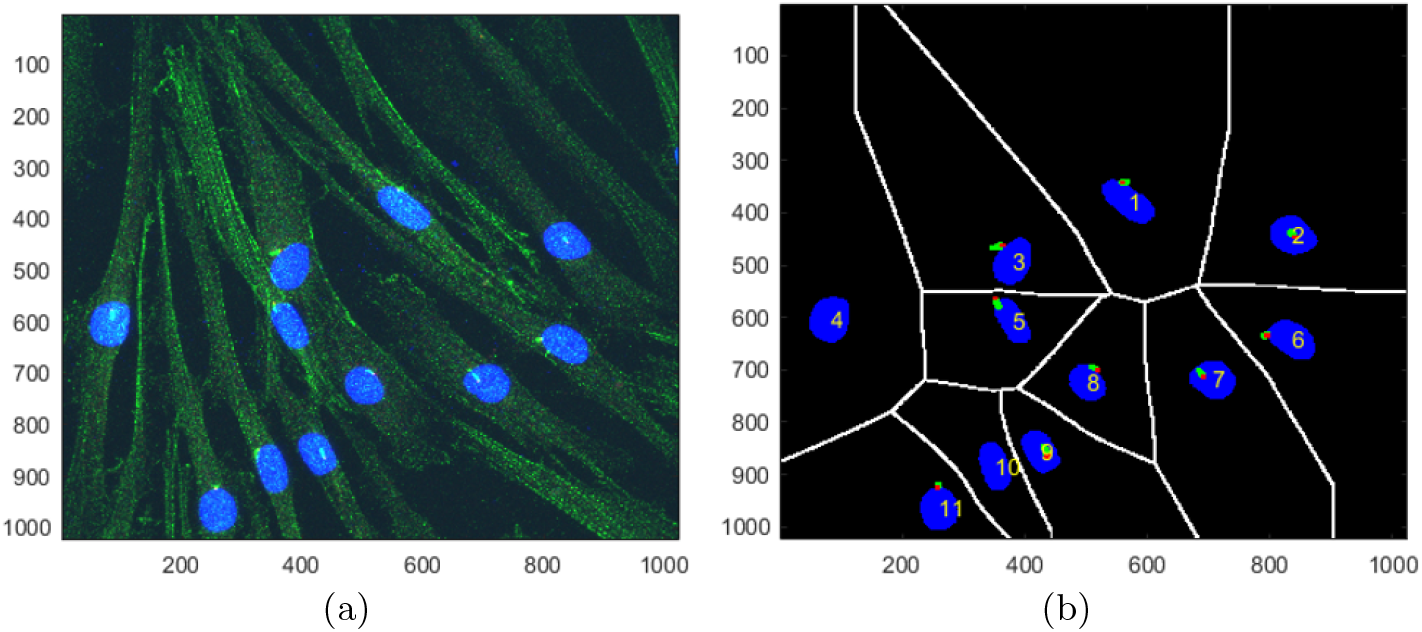
Illustration of the outcome of the segmentation pipeline. (a) One representative image with 11 cells. (b) Cells with nuclei and cilia segmented and their individual regions indicated with white lines.

**Fig. 5.**
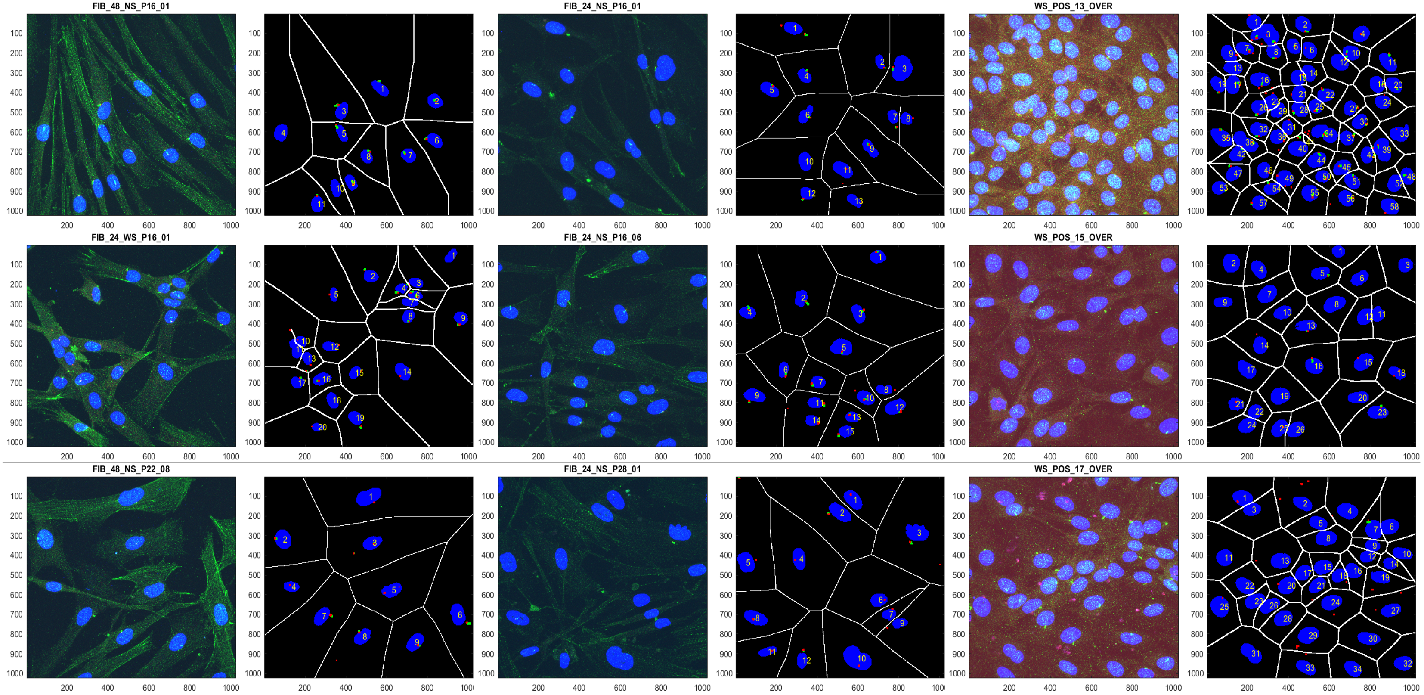
Illustration of the robustness of the segmentation process with nine images with different intensity characteristics and a varied number of cells. It should be noticed the accurate segmentation in all cases, especially in the particular dense cases on the right column.

#### Extraction of measurements

A number of metrics were extracted from the segmented images. Namely, number of nuclei, cilia axonemes and basal bodies, dimensions of nuclei, cilia axonemes and basal bodies (Fig. 6). From these, the ciliation frequency was calculated as the ratio of number of cilia to number of nuclei.

**Fig. 6.**
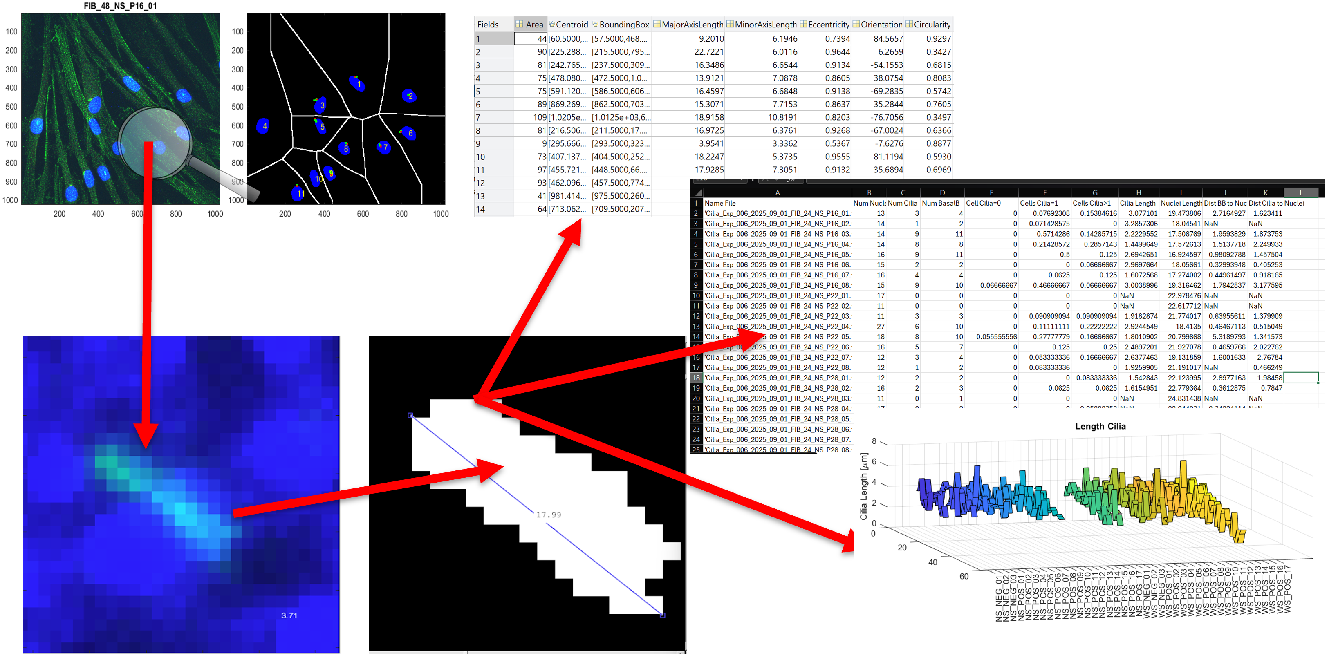
Illustration of the automated process of extraction of metrics. Cells were assessed individually. For each cell, a number of metrics were collected from the cilia, nuclei and basal body.

#### Validation of measurements

Two main metrics were validated for statistical comparison in subsequent steps: ciliation frequency and cilia length. Ciliation frequency was easily validated by a visual inspection of a number of representative cases. The length of the cilia was validated by comparing the measurements obtained with the automated methodology against manually measured lengths (Fig. 7). Manual distance measurements were obtained with ImageJ [2] without knowledge of the automated measurements.

**Fig. 7.**
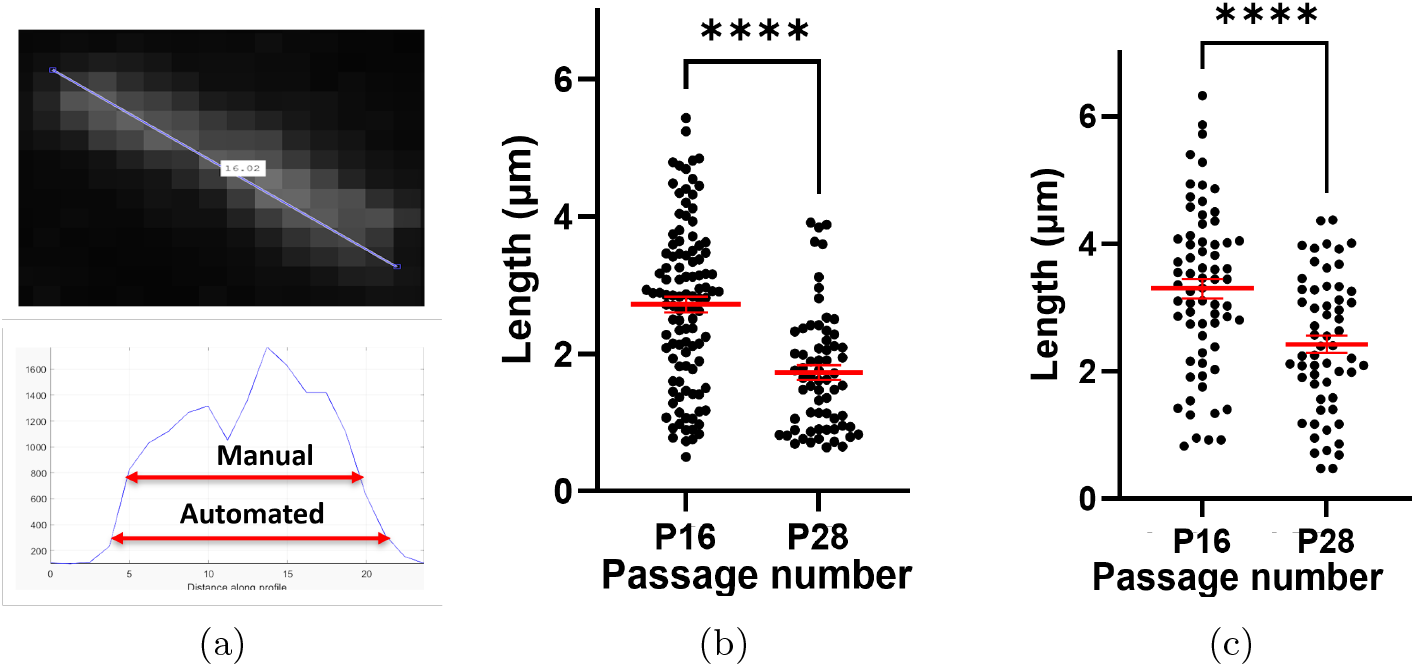
Validation of the cilia length. (a) Manual measurement of cilia length was detected to be consistently smaller than the automatically detected length. After investigation, it was noticed that the difference was due to the visual estimation of the length, which could be determined at different values of the intensity as indicated in the profile of the corresponding cell intensity. (b,c) Statistical comparison of cilia length in cells of different age (passage numbers) with (b) manual measurements and (c) automatic measurements. In both cases, the cilia shortened with age and the lengths were statistically different validating the automated process.

## 3 Results and Discussion

In total, 99 images were analysed and 1,384 individual cells were segmented. We found that the automated approach reached the same conclusion as the manual analysis (Fig. 7b and c). In both approaches, there was a statistical difference after two-tailed t-tests (**Manual measurements** mean ± SEM P16 (n=108) 2.72 ± 0.11; P28 (n=66) 1.73 ± 0.11; p < 0.0001, **Automated measurements** mean ± SEM P16 (n=68) 3.3 ± 0.16; P28 (n=59) 2.42 ± 0.14; p < 0.0001). When the manual and automated measurement values were compared, however, it was revealed that the manual measurements were consistently lower, by approximately 15% than those produced by the automated approach. Further analysis indicated that this discrepancy mainly arose from differences in the placement of measurement endpoints in manual annotations. That is, when a distance is manually measured over an intensity image, the intensity decreases from the maximum values to the minimum values which correspond to the background intensity. Thus, the decision about where to locate the end points is subjective, and in this case it was higher in intensity compared to automated measurements (Fig. 7a).

To evaluate whether changes in ciliary morphology provide a robust and quantifiable ageing-related signal, we analysed the relationship between measured cilia length and cell passage number in culture, using passage number as an operational proxy for progressive cellular ageing. When the cilia axoneme length measurements at P22 and P28 were compared to those of P16 (Fig. 8a), the results shown as a fold change are P16 =1 ± 0.05 (n=93), P22 = 0.99 ± 0.05 (n=78) and P28 = 0.70 ± 0.05 (n=63). One way ANOVA with Tukey’s multiple comparison test indicated a statistically significant difference between the groups; P16 vs P28 (p ≤0.0001); P22 vs P28 (p=0.0003). We next investigated the ciliation frequency between the groups. The results show that the proportion of ciliated cells are 96/117 (82.1%) for P16; 70/96 (72.92%) for P22; 65/97 (67.01%) for P28 (Fig. 8b). These results demonstrate a clear reduction in both cilia length and ciliation frequency with increasing cell passage number.

**Fig. 8.**
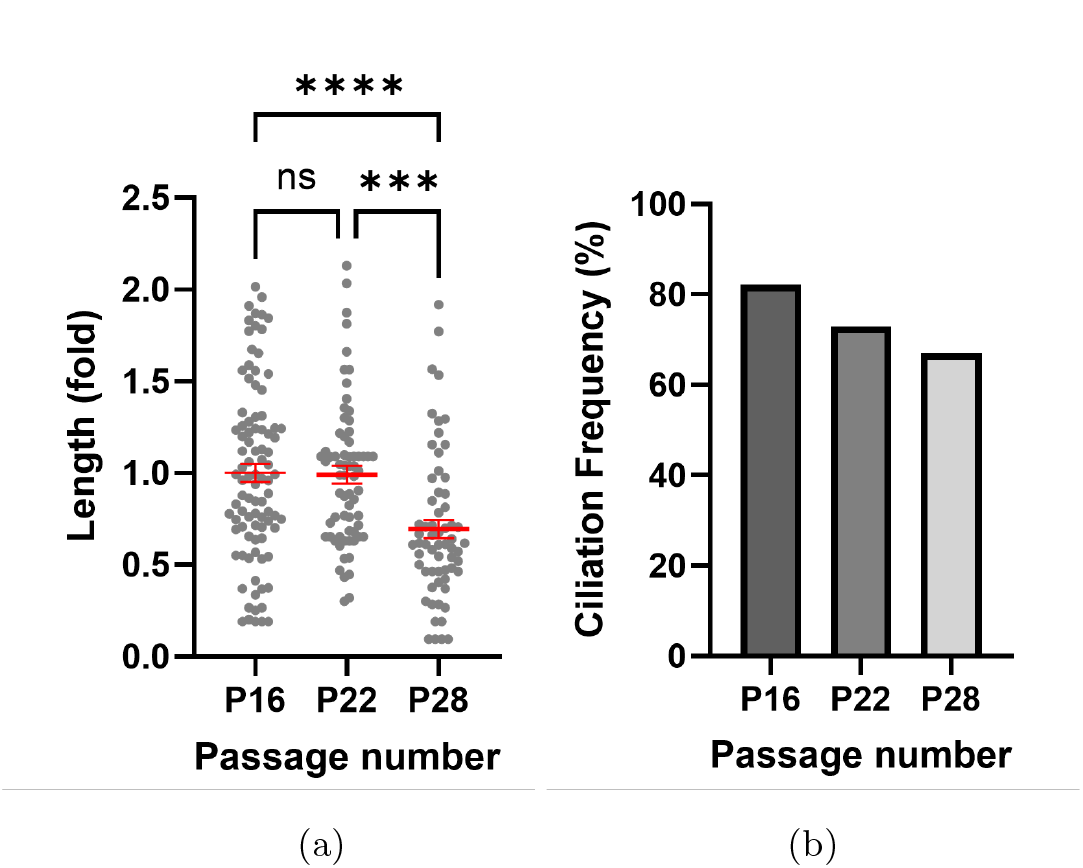
Cilia length and ciliation frequency decrease with the passage numbers in culture, representing the age of cells in vitro. The length of the cilia and the ciliation frequency (percentage of cells that contained cilia) both decreased as the number of passages increased, suggesting that the age of a cell is linked to these two metrics. This phenomenon indicates that the metabolism and maintenance of cilia decreases during vitro ageing process, thus suggesting that our approach can be used as a biomarker of ageing.

Although these findings were derived from *in vitro* experiments, the results indicate that both metrics exhibit a clear and systematic association with cellular ageing. While validation using *in vivo* data is required to assess broader physiological relevance, the proposed methodology is directly extensible to such datasets without modification. Overall, this study demonstrates that the presented framework can reliably extract cilia-derived quantitative features that serve as informative biomarkers for assessing age-related cellular changes.

## 4 Conclusions

Automatic analysis of cilia-related datasets, powered by a combination of artificial intelligence and computational algorithms can provide reliable and reproducible results, while significantly reducing analysis time and enabling high-throughput data processing. It will also aid validation of ciliation status as a potential new biomarker for health and ageing, which can be used to predict the risk of ageing-associated diseases. Application of this work in clinical context, focused on ageing-associated metabolic conditions such as nonalcoholic steatohepatitis (NASH) and sarcopenia (age-related muscle loss) is in progress.

## Acknowlegements

This work was partially funded by the National Research Foundation Korea for the project RS-2024-00509145 *Global Joint Research Center for the Development of Innovative Technology for Controlling Aging Based on Primary Cilia Metabolism*.

